# Combining Semantic Similarity and GO Enrichment for Computation of Functional Similarity

**DOI:** 10.1101/155689

**Authors:** Wenting Liu, Jianjun Liu, Jagath C. Rajapakse

**Author notes:** Emails: Wenting Liu, Jianjun Liu, Jagath C. Rajapakse.

## Abstract

Functional similarity between genes is widely used in many bioinformatics applications including detecting molecular pathways, finding co-expressed genes, predicting protein-protein interactions, and prioritization of candidate genes. Methods evaluating functional similarity of genes are mostly based on semantic similarity of gene ontology (GO) terms. Though there are hundreds of functional similarity measures available in the literature, none of them considers the enrichment of the GO terms by the querying gene pair. We propose a novel method to incorporate GO enrichment into the existing functional similarity measures. Our experiments show that the inclusion of gene enrichment significantly improves the performance of 44 widely used functional similarity measures, especially in the prediction of sequence homologies, gene expression correlations, and protein-protein interactions.

**Software availability:** The software (python code) and all the benchmark datasets evaluation (R script) are available at https://gitlab.com/liuwt/EnrichFunSim.

## Background

With the advancement of high-throughput experimental techniques, omics data are increasingly being gathered and understanding biological knowledge embedded therein requires standard and controlled organization of biological vocabularies or ontologies that represent abstract descriptions of domain-specific knowledge. Gene ontology (GO) provides a controlled vocabulary arranged in a hierarchy of terms, and facilitates annotation of gene functions and molecular attributes. Sematic similarity quantitatively measures the relationships between two terms of GO and is widely used in deriving functional similarity between two genes. Functional similarity measure is widely used in inferring genetic interactions, functional interactions, protein-protein interactions (Pesquita et al. 2008), biological pathways (Bien et al. 2012) (Guo et al. 2006), priorities of candidate genes (Moreau & Tranchevent 2012), and disease similarities (Cheng et al. 2014).

Semantic similarity measures the similarity of two ontology terms by typically evaluating their commonness normalized to their uniqueness in terms of information contents (Harispe et al. 2014)(Ranwez et al. 2014). The commonness of two terms is typically evaluated by the information content of the lowest/closest common ancestor as used by Resnik(Resnik 1999), Lin(Lin 1998), Nunivers(Mazandu & Mulder 2013), relevance similarity(Schlicker et al. 2006) measures; or by the information content of all common ancestors as evaluated by XGraSM(Couto & Silva 2011) and TopoICSim (Ehsani & Drabløs 2016). The uniqueness of GO terms are often evaluated by taking the average of the information content (IC) of the two terms. The IC of a term depends on that of the annotating corpus (Mazandu & Mulder 2014); and the topological position or semantic distance of the terms is based using ontology hierarchy as evaluated us SORA (Teng et al. 2013); or a combination of both (Wu et al. 2013).

Functional similarity (*funsim*) between two genes is typically derived using various combinations of semantic similarities between GO terms annotated to the two genes, such as the average (AVG)(Lord et al. 2003), maximum (MAX)(Mato et al. 2005), average best-matches (ABM)(Mazandu & Mulder 2013), or best-match average (BMA)(Mazandu & Mulder 2014). Alternatively, several variants of AVG and MAX combinations have also been proposed: for example, SORA(Teng et al. 2013) estimates functional similarity between two genes by computing the average of IC overlap ratio of the annotating term sets; Chabalier et al. (Chabalier et al. 2007) constructed a weighted term vector where the weight measures the representativeness of the term and computed the semantic similarity between gene products without considering their hierarchical relations; and Pandey et al.(Pandey et al. 2008) proposed a statistically motivated functional similarity measure taking into account functional specificity as well as the distribution of functional attributes across entity groups.

Functional similarity measures are derived from semantic similarities depending on the ICs of annotating terms, which are estimated by assuming a uniform distribution of terms in the background corpus. This ignores the local context and the representativeness of the terms of the gene pair, which reduces the context specificity of the similarity measure. For example, the terms annotated by both genes need to be treated more importantly than when a term is annotated by one gene. To overcome this drawback of existing functional similarity measures, we propose to introduce the probability of a term annotated to a gene by incorporating GO-enrichment of the gene pair in the computation of IC of a GO term. Specifically, in the context of two genes, the probability of a GO term annotated to a gene is defined as the joint probability of the background probability and the GO enrichment of the terms annotating the two genes. Existing functional similarity (*funsim*) measures are enriched as *funsim*\* measures with this modification that includes both the GO-enrichment and GO semantic similarity in the computation of functional similarity. We demonstrate the performance of new *funsim*\* measures on 44 *funsim* measures earlier summarized by Mazandu & Mulder(Mazandu & Mulder 2014).

## Results

### Overall Performance on all datasets

We assessed *funsim*\* measures on benchmark datasets for predicting sequence similarities, gene expression (GE) correlations, and protein-protein interactions (PPI) and compared with those of *funsim* measures. Table 1 shows one-sided *p*-values on the improvement of performances, using Wilcoxon signed rank tests(Wilcoxon 1945) of all the experiments on four benchmark datasets. As seen, *funsim*\* measures showed a significant improvement over *funsim* measures in the prediction of protein interactions on 132 experiments on yeast PPI data, gene co-expressions on 132 experiments using yeast GE data, and sequence similarities on 264 experiments on CESSM dataset; and on all 528 experiments. Irrespective of the ontology (BP, MF, or CC) and the type of *funsim* measure, the incorporation of GO enrichment in *funsim*\* measure significantly improved the prediction of sequence similarities, gene co-expression patterns, and protein-protein interactions.

**Table 1.** The details of three datasets and statistical significances of the improvement of performances of *funsim*\* over *funsim* measure: yeast PPI dataset, yeast GE dataset, and sequence similarities (ECC, Pfam, and SeqSim) on protein pairs given by CESSM.

| DataType | DataSets | #protein pairs | ontology | #experiments | *p*-value |
| --- | --- | --- | --- | --- | --- |
| yeast_PPI | PPI_BP; PPI_MF; PPI_CC | 6000 | BP | 132 | 2.03E-05 |
| yeast_GE | GE_BP; GE_MF; GE_CC | 4800 | BP; MF; CC | 132 | 9.90E-08 |
| CESSM | ECC; Pfam; SeqSim | 13430 | BP; MF | 264 | 3.48E-15 |
| Total | PPI; GE; ECC; Pfam; SeqSim | 45830 | BP; MF; CC | 528 | < 2.2e-16 |

### Performance of funsim* measures on three types of biological data

Table 2 lists 10 top performed *funsim* measures, the corresponding *funsim*\* measures, percentages of performance improvement of *funsim*\* over *funsim*, and statistical significances of the improvements on different datasets. The significances of improvement were computed using Williams test (DA Williams 1972) (FDR adjusted *p*-values). The list of 44 *funsim* measures is given in Table 3. Among the 44 *funsim* measures, *funsim*\* improved for the top performers on almost all of them.

**Table 2.** Performances of top 10 *funsim* measures, and corresponding *funsim*\* values, percentage improvement of *funsim*\* over *funsim*, and statistical significance of improvement on each dataset.

| DataSets | Methods | *funsim* | *funsim* * | Improvement(%) | FDR p-value |
| --- | --- | --- | --- | --- | --- |
| ECC_BP | XNBMA | **0.4748** | **0.4748** | **0.00** | 3.99E-01 |
|  | XLBMA | **0.4708** | **0.4748** | **0.84** | **2.53E-253** |
|  | NBMA | 0.4635 | 0.4651 | 0.33 | **2.25E-51** |
|  | XNABM | **0.4600** | **0.4605** | **0.10** | **4.47E-05** |
|  | WDIC | **0.4553** | **0.4554** | **0.04** | **9.96E-02** |
|  | XLABM | 0.4547 | 0.4556 | 0.21 | **1.31E-10** |
|  | SBMA | 0.4511 | 0.4523 | 0.26 | **4.89E-16** |
|  | LBMA | 0.4497 | 0.4518 | 0.46 | **6.49E-64** |
|  | ZDIC | 0.4490 | 0.4490 | 0.02 | 3.99E-01 |
|  | ZBMA | 0.4472 | 0.4491 | 0.41 | **5.23E-50** |
| Pfam_BP | WBMA | 0.4716 | **0.5223** | **10.74** | **1.64E-44** |
|  | WABM | 0.4621 | **0.5261** | **13.84** | **1.50E-71** |
|  | UBMA | **0.4764** | **0.4764** | **0.00** | 3.99E-01 |
|  | UABM | **0.4754** | **0.4758** | **0.08** | 2.41E-01 |
|  | XNBMA | 0.4721 | 0.4752 | 0.64 | **3.37E-209** |
|  | XNABM | 0.4673 | 0.4710 | 0.78 | **1.25E-236** |
|  | XLBMA | 0.4590 | 0.4752 | **3.53** | **0.00E+00** |
|  | ZGIC | **0.4665** | **0.4670** | **0.11** | **4.94E-32** |
|  | ZDIC | **0.4606** | **0.4616** | **0.22** | **9.12E-89** |
|  | AGIC | 0.4607 | 0.4608 | 0.04 | **4.19E-02** |
| SeqSim_BP | AGIC | **0.7622** | **0.7633** | **0.14** | **1.21E-248** |
|  | ZGIC | 0.7592 | 0.7603 | 0.15 | **7.71E-293** |
|  | UGIC | 0.7584 | 0.7570 | **-0.19** | **2.98E-06\*** |
|  | WGIC | 0.7446 | 0.7450 | 0.06 | **1.40E-29** |
|  | XNBMA | **0.7256** | **0.7283** | **0.37** | **2.10E-263** |
|  | UDIC | **0.7281** | **0.7247** | **-0.46** | **1.92E-20\*** |
|  | UUIC | **0.7368** | **0.7126** | **-3.28** | **3.53E-72\*** |
|  | XNABM | **0.7227** | **0.7262** | **0.49** | **0.00E+00** |
|  | ADIC | **0.7228** | **0.7246** | **0.24** | **0.00E+00** |
|  | XLBMA | 0.7091 | 0.7283 | **2.71** | **0.00E+00** |
| ECC_MF | XNBMA | **0.7525** | **0.7567** | **0.55** | **0.00E+00** |
|  | NBMA | 0.7485 | 0.7525 | 0.54 | **7.68E-226** |
|  | XLBMA | 0.7362 | 0.7421 | 0.80 | **0.00E+00** |
|  | WBMA | 0.7100 | **0.7665** | **7.96** | **2.59E-157** |
|  | **XNABM** | **0.7292** | **0.7337** | **0.61** | **0.00E+00** |
|  | NABM | 0.7248 | 0.7294 | 0.64 | **1.26E-243** |
|  | **SBMA** | 0.7189 | 0.7229 | 0.56 | **1.50E-75** |
|  | LBMA | 0.7176 | 0.7238 | 0.87 | **1.58E-260** |
|  | WABM | 0.6889 | **0.7479** | **8.56** | **2.13E-152** |
|  | ZBMA | 0.7145 | 0.7171 | 0.36 | **1.40E-90** |
| Pfam_MF | **AGIC** | **0.6170** | **0.6203** | **0.53** | **2.08E-124** |
|  | **XNABM** | **0.5829** | **0.5849** | **0.34** | **5.17E-49** |
|  | XNBMA | **0.5818** | **0.5833** | **0.27** | **3.68E-30** |
|  | **ADIC** | **0.5710** | **0.5769** | **1.03** | **0.00E+00** |
|  | AUIC | **0.5729** | **0.5729** | **0.00** | 3.99E-01 |
|  | XLBMA | 0.5655 | 0.5673 | 0.32 | **3.34E-21** |
|  | XLABM | 0.5650 | 0.5673 | 0.42 | **1.16E-31** |
|  | WABM | **0.5034** | **0.6283** | **24.82** | **0.00E+00** |
|  | WBMA | 0.5003 | **0.6260** | **25.11** | **0.00E+00** |
|  | NABM | 0.5259 | 0.5309 | 0.95 | **2.69E-185** |
| SeqSim_MF | **AGIC** | **0.6285** | **0.6358** | **1.17** | **0.00E+00** |
|  | AUIC | **0.5510** | **0.5510** | **0.00** | 3.99E-01 |
|  | **ADIC** | **0.5288** | **0.5375** | **1.64** | **0.00E+00** |
|  | ZGIC | 0.5127 | 0.5146 | 0.36 | **1.45E-255** |
|  | **XNABM** | **0.5049** | **0.5058** | **0.18** | **1.51E-09** |
|  | XNBMA | **0.5017** | **0.5021** | **0.08** | **7.50E-03** |
|  | XLABM | 0.4922 | 0.4931 | 0.18 | **3.25E-05** |
|  | **WAVG** | **0.4313** | **0.5514** | **27.84** | **1.42E-184** |
|  | XLBMA | 0.4910 | 0.4912 | 0.04 | 2.70E-01 |
|  | WABM | 0.4050 | **0.5425** | **33.95** | **0.00E+00** |
|  | WBMA | 0.4001 | **0.5370** | **34.21** | **0.00E+00** |
| PPI_BP | XNBMA | **0.8785** | **0.8791** | **0.07** | **3.19E-10** |
|  | XNMAX | 0.8780 | 0.8784 | 0.04 | **3.84E-06** |
|  | XLMAX | **0.8778** | **0.8781** | **0.04** | **3.69E-03** |
|  | SMAX | 0.8777 | 0.8777 | -0.01 | 3.99E-01 |
|  | LMAX | 0.8774 | 0.8776 | 0.03 | **9.64E-02** |
|  | NMAX | 0.8767 | 0.8770 | 0.03 | **6.30E-03** |
|  | XLBMA | 0.8758 | 0.8765 | 0.08 | **7.45E-11** |
|  | ZMAX | 0.8742 | 0.8743 | 0.01 | 2.71E-01 |
|  | **XNABM** | **0.8737** | **0.8746** | **0.11** | **1.52E-19** |
|  | NBMA | 0.8726 | 0.8735 | 0.10 | **4.52E-19** |
| PPI_MF | UBMA | **0.7452** | **0.7454** | **0.03** | 2.00E-01 |
|  | **ADIC** | **0.7442** | **0.7456** | **0.19** | **6.67E-28** |
|  | **AGIC** | **0.7442** | **0.7456** | **0.19** | **3.37E-35** |
|  | AUIC | **0.7440** | **0.7448** | **0.10** | 3.86E-01 |
|  | ZGIC | 0.7429 | 0.7428 | -0.01 | 2.50E-01 |
|  | ZDIC | 0.7429 | 0.7428 | -0.01 | 2.90E-01 |
|  | UABM | **0.7422** | **0.7426** | **0.06** | **1.15E-02** |
|  | ZUIC | 0.7394 | 0.7403 | 0.12 | 3.85E-01 |
|  | NBMA | 0.7383 | 0.7403 | 0.27 | **3.90E-13** |
|  | UMAX | **0.7389** | **0.7389** | **-0.01** | 3.99E-01 |
| PPI_CC | UABM | **0.8217** | **0.8237** | **0.25** | **6.21E-04** |
|  | ZBMA | **0.8219** | **0.8228** | **0.11** | **1.58E-06** |
|  | ZABM | 0.8202 | 0.8213 | 0.13 | **2.05E-09** |
|  | NABM | 0.8187 | 0.8197 | 0.13 | **5.02E-11** |
|  | UBMA | 0.8180 | 0.8203 | **0.28** | **1.11E-04** |
|  | NBMA | 0.8186 | 0.8195 | 0.11 | **4.13E-09** |
|  | XNABM | 0.8176 | 0.8181 | 0.06 | **3.09E-05** |
|  | SBMA | 0.8164 | 0.8153 | **-0.14** | **2.48E-07\*** |
|  | LBMA | 0.8143 | 0.8154 | 0.13 | **8.88E-10** |
|  | XLABM | 0.8144 | 0.8150 | 0.07 | **1.46E-05** |
| GE_BP | AGIC | **0.2873** | **0.2877** | 0.14 | **6.90E-03** |
|  | ZGIC | 0.2869 | 0.2875 | 0.20 | **2.45E-06** |
|  | WGIC | 0.2839 | 0.2843 | 0.15 | **8.14E-03** |
|  | UABM | **0.2826** | **0.2855** | **1.00** | **1.05E-02** |
|  | ADIC | 0.2829 | 0.2839 | 0.35 | **2.20E-09** |
|  | UDIC | **0.2812** | **0.2854** | **1.50** | **6.73E-04** |
|  | UBMA | **0.2800** | **0.2828** | **0.99** | **1.25E-02** |
|  | AUIC | **0.2851** | **0.2774** | **-2.69** | **5.28E-02\*** |
|  | ZDIC | 0.2798 | 0.2809 | 0.40 | **1.32E-13** |
|  | UGIC | **0.2762** | **0.2815** | **1.93** | **4.43E-06** |
| GE_MF | AGIC | **0.2022** | **0.2023** | 0.05 | 3.69E-01 |
|  | ADIC | **0.2002** | **0.2008** | **0.26** | **1.84E-02** |
|  | AUIC | 0.1957 | 0.1973 | 0.81 | 3.69E-01 |
|  | XNBMA | 0.1905 | 0.1921 | **0.85** | **6.18E-03** |
|  | WGIC | 0.1898 | 0.1894 | **-0.21** | **6.39E-03\*** |
|  | UABM | 0.1886 | 0.1902 | **0.88** | **9.67E-07** |
|  | XNABM | 0.1886 | 0.1902 | **0.88** | **6.69E-03** |
|  | UBMA | 0.1886 | 0.1902 | **0.85** | **9.67E-07** |
|  | ZGIC | 0.1885 | 0.1881 | **-0.20** | **2.24E-02\*** |
|  | ZDIC | 0.1839 | 0.1838 | -0.06 | 3.68E-01 |
| GE_CC | ZDIC | **0.4253** | **0.4263** | **0.24** | **8.32E-09** |
|  | ZGIC | **0.4233** | **0.4236** | **0.07** | **5.59E-02** |
|  | ADIC | 0.4220 | 0.4229 | 0.21 | **2.53E-07** |
|  | ZUIC | **0.4229** | **0.4189** | -0.95 | 2.89E-01 |
|  | AGIC | 0.4202 | 0.4204 | 0.05 | 1.89E-01 |
|  | AUIC | 0.4190 | 0.4158 | -0.76 | 3.12E-01 |
|  | WGIC | 0.4081 | 0.4077 | -0.10 | **8.64E-04** |
|  | WDIC | 0.4063 | 0.4064 | 0.04 | 3.12E-01 |
|  | WUIC | 0.4037 | 0.3932 | **-2.59** | **4.92E-02\*** |
|  | UABM | 0.3940 | 0.3997 | **1.45** | **1.24E-06** |
|  | UDIC | 0.3920 | 0.3980 | **1.55** | **9.21E-10** |

**Supplementary Tables** 4-7 give the details of performances of *funsim*\* over all 44 *funsim* measures in predicting sequence similarities, gene co-expressions, and protein-protein interactions. **Supplementary Figures** 1-4 show the improvement of evaluation scores from *funsim* to *funsim*\* on sequence homologies on BP ontology and MF ontology, gene co-expression correlations, and PPIs on three ontologies, respectively.

Our experiments on different *funsim* measures yielded similar observations as seen by Mazandu & Mulder(Mazandu & Mulder 2014). In general, BMA and ABM methods provide the best performances and performed equally well on most semantic similarity measures. Adaptation of efficient correction factors improved the performance on some measures: Schlicker(Schlicker et al. 2006) uses the IC value of MICA and does not significantly improve the performance of the Lin(Lin 1998) approach; XGraSM(Couto & Silva 2011) uses all common informative ancestors to correct Lin(Lin 1998) and Nunivers(Mazandu & Mulder 2013) approaches in order to improve their performances. Thus, including common informative ancestors in the conception of a semantic similarity improves its performance, especially for approaches that include only the features of child terms in the computation of IC. This is the case for the annotation-based Zhang(Zhang et al. 2006) and Wang(Wang et al. 2007) approaches, where the SimGIC(Pesquita et al. 2008) measure shows the overall best performance.

Lin(Lin 1998), Nunivers(Mazandu & Mulder 2013), GO-universal(Mazandu & Mulder 2013), Wang(Wang et al. 2007), and SimGIC(Pesquita et al. 2008) measures improved much more significantly than other measures with the incorporation of GO enrichment. As the *funsim*\* measure differently treats unique GO terms (annotated to only one gene) and common terms annotated by two genes, measures consisting of both kinds of terms are significantly improved with GO enrichment: for example, Lin(Lin 1998), Nunivers(Mazandu & Mulder 2013), and SimGIC(Pesquita et al. 2008) measures consider both common terms and individual terms; GO- universal(Mazandu & Mulder 2013) measure considers all children terms (common or individual terms); and Wang(Wang et al. 2007) measure consider all ancestors (common terms) and children terms (common or individual terms). Especially, Wang(Wang et al. 2007) measure (WABM, WBMA) improved significantly on capturing sequence homology, with an correlation improvement of 8% of ECC, 25% of Pfam, 34% of SeqSim on MF ontology; and 13% of Pfam, 16% of SeqSim on BP ontology; GO-universal approach (UABM, UBMA) improved most significantly (labelled as green) for GE correlations on three ontologies, and inferring PPIs on CC; XGraSM of Nunivers approach (XNABM, XNBMA) improved most significantly for GE correlations on MF, and inferring PPIs on BP; and annotation-based SimGIC and SimDIC are improved most significantly for inferring PPIs on MF. Out of all 44 *funsim* measures, the performance of measure related to UIC measure didn’t improve with GO enriched *funsim*\* measures. This is because the UIC measure does not discriminate common terms and unique terms while the enrichment is manifested by the differences between common and unique terms.

## Conclusions

From GO annotations, many *funsim* measures have been proposed for efficient exploitation of biological knowledge embedded in omics data. These measures were derived based on the topological structure of GO semantics and GO annotations of the genes/proteins annotating (background) corpus. However, the representativeness of GO terms of two querying genes has been neglected in deriving their functional measures. We proposed an enriched functional similarity between two genes, *funsim*\*, that incorporates the enrichment of GO terms of the genes and demonstrated improvements of performance of a large majority of *funsim* measures in the literature.

We tested *funsim*\* measures on 44 *funsim* measures on three benchmark datasets including sequence similarities given by the CESSM dataset, yeast GE data, and yeast PPI data. We performed a quantitative performance evaluation of *funsim* measures that adopt different methods for evaluating IC and combining semantic similarities of GO terms. Results indicate that *funsim*\* generally improves the performance of *funsim* measures in predicting sequence similarities, gene co-expressions, and protein-protein interactions. We conclude that the enrichment by the querying genes is a necessary step when computing their functional similarities. Especially, for *funsim* measures considering both common terms and individual terms of the two genes, e.g., Wang approach, the performances of *funsim*\* improved much significantly over *funsim* measure. We also noticed that *funsim*\* significantly improved the performance especially on datasets containing a lot of uniquely annotated genes (i.e., those in the low levels of GO hierarchy).

*Funsim*\* is easily adapted to and generally improves the performance of any *funsim* measure. One could extend our method to evaluate the functional coherence of gene sets, which will have applications in the detection of functional modules or pathways. On the other hand, the accuracy of GO annotation naturally limits the performance of existing *funsim* measures as they do not consider both the local context of two genes and the background distribution of terms in the annotating corpus. Our experiments suggest that the local context of querying genes is sensitive to the missing and spurious terms in the GO annotating corpus. *Funsim*\* measures help find the most significant functionally similar genes and provide more reliable computational evidences for finding new pathways and disease genes. We conclude that the GO enrichment is an essential step when assessing functional similarity of two genes.

## Online Methods

### Data Sets

We investigated the performance of *funsim*\* by evaluating their correlations with sequence similarities, gene co-expressions, and protein-protein interactions. Molecules with sequence similarities are likely to have similar functions or MF ontology. Molecules with similar gene expressions are likely to belong to the same pathway or have similar BP ontology. Interacting proteins are located in the same cellular location, so likely to have the similar CC ontology. We adopted the same benchmark datasets used by earlier comprehensive studies evaluating *funsim* measures. Correlations of *funsim* and enriched *funsim*\* measures with protein sequence similarities from Collaborative Evaluation of Semantic Similarity Measures (CESSM) online tool (Pesquita et al. 2009), gene expression (GE) correlations (Yang et al. 2012) and AUC scores on predicting protein-protein interactions(Pesaranghader et al. 2015) were evaluated. Experimental results demonstrate that the enriched functional similarity measure *funsim*\* significantly improves the performance over existing *funsim* measures on benchmark datasets.

### Correlation with sequence similarity

Various studies have shown that similar sequences have similar ontological annotations(Lord et al. 2003) and used sequence similarities to demonstrate the goodness of similarity measures (Yang et al. 2012) (Pesaranghader et al. 2015). For BP and MF, we use the CESSM online tool (Pesquita et al. 2009) (http://xldb.di.fc.ul.pt/tools/cessm/) and downloaded the dataset of selected human proteins with known relationships to compare different measures. The CESSM website provides a list of protein pairs and similarity between pairs of proteins, using three distinct evaluations: sequence similarity (SeqSim), Pfam domain similarity, and enzyme commission class (ECC) similarity. High correlations between protein similarities captured by SeqSim, Pfam similarity, and ECC similarity indicate the goodness and unbiasedness of a *funsim* measure.

### Correlation with gene expressions

Genes involved in the same biological process, sharing similar functions or cellular components, tend to exhibit similar expression patterns, so a good correlation should exist between co-expressed genes and functional similarities. We used the same gene-expression dataset of *S. cerevisiae*, used by earlier studies (Yang et al. 2012) (Pesaranghader et al. 2015), which contains co-expression values of 4800 pairs of genes for each ontology, downloaded from GeneMANIA (Gillis & Pavlidis 2013) and other microarray experiments. We computed Pearson’s correlations between gene co-expressions and functional similarity values of BP, MF and CC ontologies.

### AUC on predicting protein-protein interactions

Two interacting proteins have the same CC, share similar functions, and are likely to belong to same BP. Therefore, functional similarity between two proteins is an indicative of an interaction (Chabalier et al. 2007) (Maetschke et al. 2012). Similar to an earlier study (Pesaranghader et al. 2015), we formulated the prediction of protein-protein interaction (PPI) as a classification problem using functional similarities of the two proteins. Above a certain threshold of functional similarity, an interaction is identified between two proteins. We gathered data from a yeast dataset (Pesaranghader et al. 2015) containing 6,000 PPI pairs for each gene ontology where about half of the data are positive interactions from a core subset of the Database of Interacting Proteins (DIP) (Salwinski et al. 2004); and the other half are negative interactions generated by randomly choosing annotated protein pairs in that ontology. For evaluation, we used the area under the curve (AUC) values of the receiver operating characteristic (ROC) curve of the predictor. The ROC curves plot the true positive rate (sensitivity) vs false positive rate (1-specificity) values for prediction at different thresholds.

### Significance test for correlation improvement

To show any improvement of the enriched *funsim* measures, *funsim*,* we computed the improved percentage of the correlations between *funsim* score and sequence similarity score and gene co-expression score, and of AUC of prediction of PPIs. To determine the statistical significance of an improvement of correlation or AUC values for each *funsim* to *funsim*\* measure, we adopted Williams test (DA Williams 1972) for correlations between two metrics (Steiger. 1980) (Graham & Baldwin 2014). Specifically, to test whether the population correlation between *X*_1_ and *X*_3_ equals the population correlation between *X*_2_ and *X*_3_, we computed the following *t*-test:

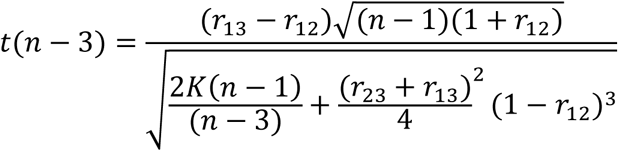

Where 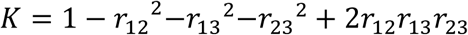.

The higher the correlation between the metric scores, the greater is the statistical power of this test than the Fisher *r* to *z-*transformation test on independent correlations. As *funsim* and *funsim*\* are highly correlated, we used this Williams test (DA Williams 1972)and adopted FDR for multiple test correction.

To determine whether correlations or AUC values are significantly improved for all *funsim* measures to *funsim*\* on each dataset (CESSM, yeast GE, yeast PPI, and the combination of the three datasets), we implemented the Wilcoxon signed rank test with continuity correction(Longnecker 1983), which tests repeated measurements on a single sample to assess whether their population mean ranks differ. This test is suggested as an alternative for *t*-test for dependent samples when the population cannot be assumed to be normally distributed. We used one-sided Wilcoxon signed rank test to show whether *funsim*\* significantly improves the performance of *funsim* irrespective of the *funsim* measure and the type of ontology.

### Funsim measures

#### The information content of a gene ontology term

Gene ontology (GO) describes an ontology of terms describing how gene products behave in a cellular context in a species-independent manner. Gene ontology covers three domains: biological process (BP), molecular function (MF), and cellular component (CC). BP is a collection of molecular events, MF defines gene functions in biological processes, and CC describes gene locations within a cell. A gene is associated with GO terms that describe the properties of its products (i.e., proteins), and the annotation corpus or gene ontology annotation (GOA) file corresponds to an organism.

There are three semantic relations between two GO terms: *is-a* is used when one GO term is a subtype of another GO term, *part-of* is used to represent part-whole relationship in the GO terms, and *regulate* is used when the occurrence of one biological process directly affects the manifestation of another process or quality(Gene & Consortium 2000). The GO terms and their relations are constructed in a hierarchical directed acyclic graph (DAG) where the three domains, BP, MF and CC, are represented as three roots at the topmost level. Nodes/terms near the root of a DAG have broader functions and are hence shared by many genes; leaf nodes/terms on the other hand convey more specific biological functions.

GOA is the process in which gene or gene products are annotated using GO terms. GOA data can be readily downloaded from the GO annotation database (http://www.geneontology.org/GO.downloads.annotations.shtml) for a species. The hierarchical structure of GO allows annotators to assign properties of genes or gene products at different levels, depending on the availability of the information about the entity. Typically, when inferring information of a gene that is annotated by some hierarchy of GO terms, more specific information on biological functions at lower levels are chosen as the inference base due to their richer information content.

The information content of a GO term *t* is defined as

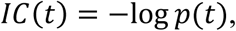

where *p*(*t*), the probability of term *t* annotating to a gene, is usually defined as the frequency of the term *t* relative to the frequency of the root term in the same ontology tree, given a corpus (e.g., an organism) of annotating genes. The term probability is given by

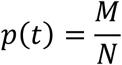

where *M* is the number of genes annotated by term *t* and *N* is the total number of genes in the annotating corpus. According to the true-path-rule, when a gene is annotated by a term, the gene should be also annotated by ancestor terms because of the hierarchical structure of GO. Thus, the frequency of the root term is equal to the number of all genes in annotating organism and this definition of *p*(*t*) assumes a uniform distribution of probabilities to randomly annotating a gene by term *t*.

In topology-based semantic measures, *p*(*t*) depends on the topological position of the term in GO-DAG. Specifically, Zhang’s method(Zhang et al. 2006) defines a D-value for a term by recursively summing gene counts of all its children from the bottom up. For a pair of terms, D-value is defined as the minimum D-value of their common ancestors. In GO-universal method (Mazandu & Mulder 2013),

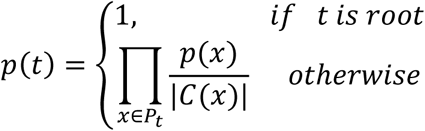

where *P_t_* is the parent term set of term *t*, and |*C*(*x*)| is the number of children with term *x* as parent.

#### Semantic similarity measures between two GO terms

Several approaches have been proposed for determining the semantic similarity measure between two GO terms, including annotation-based measures such as Resnik(Resnik 1999), Lin(Lin 1998), Jiang & Conrath(Jiang & Conrath 1997), Nunivers(Mazandu & Mulder 2013), corrections to annotation-based measures such as Graph-based Similarity (Disjunct Common Ancestor eXended GraSM, denoted as XGraSM(Couto & Silva 2011)), relevance similarity (Schlicker et al. 2006); and topology-based measures, such as Zhang(Zhang et al. 2006), GO-universal(Mazandu & Mulder 2013) and Wang(Wang et al. 2007) approaches. An implementation of these measures is provided by the A-DaGO-Fun tool(Mazandu et al. 2015).

For Resnik(Resnik 1999) measure, simantic simiarity of two terms *t*_1_ and *t*_2_ is defined as information content of their most informative common ancestor (MICA), denoted by *t*_0_.

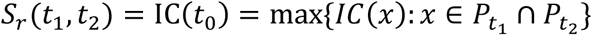

Since GO enrichment applies when both individual and common terms are considered and the Resnik(Resnik 1999) measure solely considers the terms annotated to both two genes, so both f*unsim* and *funsim*\* using Resnik measures have no difference, so Resnik measure is not considered in our assessment.

The Lin(Lin 1998) semantic similarity measure takes MICA between terms and normalized by the average of IC values of the two terms.

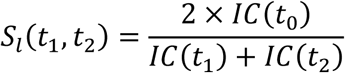

Note that the Jiang & Conrath(Jiang & Conrath 1997) measure is a particular case of Lin approach, so only Lin measure is considered in the experiments. The Nunivers(Mazandu & Mulder 2013) measure was proposed to satisfy the requirement that the similarity between a term to itself should be one:

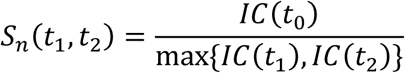

The Schlicker(Schlicker et al. 2006) measure combines Resnik(Resnik 1999) with Lin(Lin 1998) similarity as

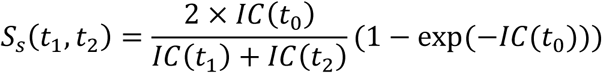

The graph-based (XGraSM(Couto & Silva 2011)) extensions of Lin(Lin 1998) and Nunivers(Mazandu & Mulder 2013) measures, respectively, are

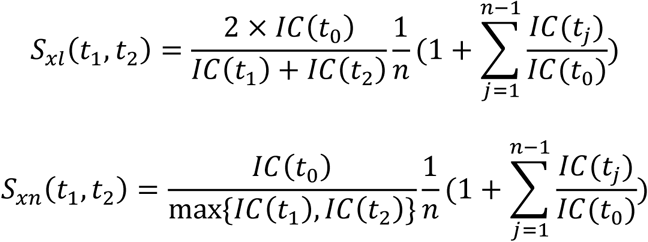

where *n* is the number of all informative common ancestors of the terms *t*_1_ and *t*_2_, the ancestor terms are ordered in an increasing order of information content, and *n*th ancestor term is MICA.

In topology-based measures by Zhang(Zhang et al. 2006), GO-universal(Mazandu & Mulder 2013) and Wang(Wang et al. 2007), the information content also incorporates the position characteristics from GO-DAG topology, and their definitions were as given in the information contents section. Wang(Wang et al. 2007) considered semantic value *s_t_* of term *t*, recursively from its children set (*C*(*x*) is the children set with term *x* as parent) based on the semantic contribution factor *w*_e_ for *is-a* and *part-of* as 0.8 and 0.6, respectively.

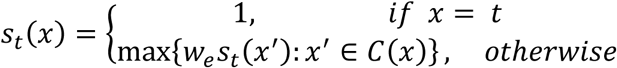

And the information content is computed from the summation of the semantic values of all its ancestors set *P_t_*,

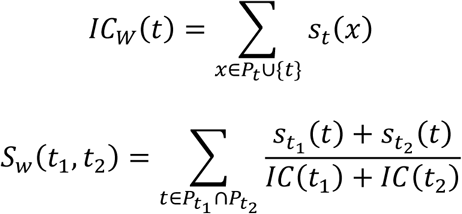

#### Functional similarity (funsim) measures between two genes

Functional similarity between two genes is computed from a combination of their annotating GO terms by using basic statistical measures of closeness (mean, max, min, etc.) such as Best-Match Average (BMA), Average Best-Matches (ABM), Average (AVG) and Maximum (MAX). These measures of closeness are known to be sensitive to biases introduced by the abnormal distances from the majority, or outliers.

*Funsim* measures based-on basic statistical measures between semantic similarities between two genes *g*_1_ and *g*_2_ are defined as

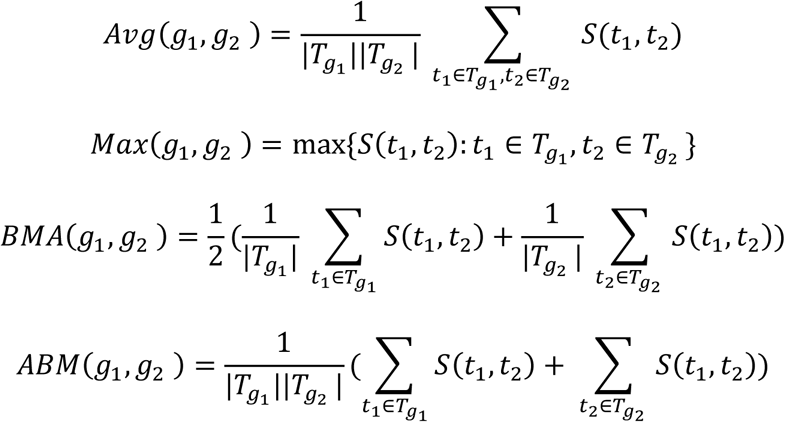

where *T_g_*_1_ is the annotated term set of gene *g*_1._

Other measures such as SimGIC(Pesquita et al. 2008), SimDIC(Mazandu & Mulder 2013), and SimUIC(Mazandu & Mulder 2013) use the IC of terms directly in the computation of functional similarity. Direct term-based *funsim* measures are defined as

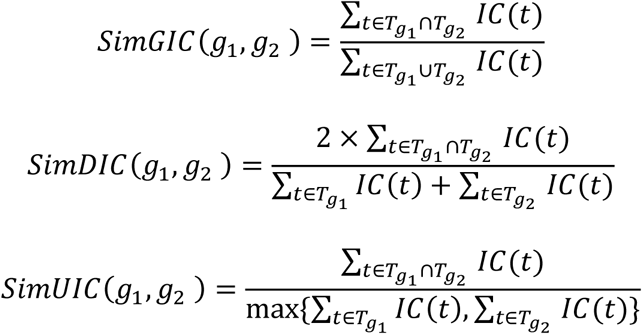

Table 3 categorizes 44 *funsim* measures in the literature, based on combination of nine GO semantic measures (the first three based on topology from GO-DAG, and the other six based on corpus annotation), basic statistical measures (MAX, AVE, BMA, ABM), and three direct term-based measures.

**Table 3.** The details and categorization of 44 *funsim* measures

| Acronyms | Semantic similarity method | Statistical Measures |  |  |  | Direct term-based Measures |  |  |
| --- | --- | --- | --- | --- | --- | --- | --- | --- |
|  |  | ABM | BMA | MAX | AVG | DIC | GIC | UIC |
| U | GO-universal (Mazandu, & Mulder., 2013) | UABM | UBMA | UMAX | UAVG | UDIC | UGIC | UUIC |
| Z | Zhang (Zhang., et al., 2006), | ZABM | ZBMA | ZMAX | ZAVG | ZDIC | ZGIC | ZUIC |
| W | Wang (Wang., 2007) | WABM | WBMA | WMAX | WAVG | WDIC | WGIC | WUIC |
| N | Nunivers (Mazandu, & Mulder., 2013) | NABM | NBMA | NMAX | NAVG | - | - | - |
| XN | XGraSM (Couto et al., 2007) on Nunivers (Mazandu, & Mulder., 2013) | XNABM | XNBMA | XNMAX | XNAVG | - | - | - |
| L | Lin (Lin., 1998) | LABM | LBMA | LMAX | LAVG | - | - | - |
| XL | XGraSM (Couto et al., 2007) on Lin (Lin., 1998) | XLABM | XLBMA | XLMAX | XLAVG | - | - | - |
| S | Relevance similarity (Schlicker, Domingues, Rahnenführer, & Lengauer, 2006) | SABM | SBMA | SMAX | SAVG | - | - | - |
| A | Annotation-based approach | - | - | - | - | ADIC | AGIC | AUIC |

#### Enriched term probability p^*^(t) and functional similarity funsim*

Earlier approaches of functional similarity (*funsim*) assume the representativeness of GO terms based on the annotating corpus (or GOA file). They fail to take into account the enrichment by the querying pair of genes. GO enrichment usually assumes a hypergeometric distribution of annotating terms *t* in a given gene set(Huang et al. 2008) and has been effectively used in finding pathways most represented by the gene set. We propose to incorporate GO-enrichment in the computation of functional similarity between a pair of genes. Specifically, for two genes, a term *t* annotated to only one gene and to both two genes should be treated differently.

The probability of term *t* annotating *k* genes by in a gene set of size *n* is given by a hypergeometric distribution as

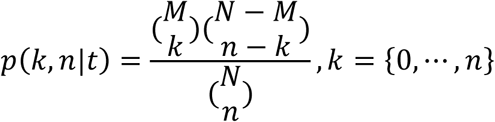

where *N* is the number of annotated genes in an organism and *M* is the number of genes annotated by term *t*. The joint probability of annotating *k* genes in a gene set with size *n* by term *t* for a corpus with *N* genes is given by

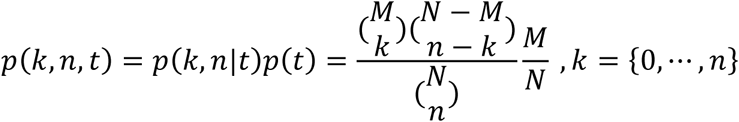

the 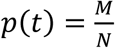 is the term probabilities inferred by the annotating corpus. We define the enriched probability term *p*\*(*t*) as this joint probability in order to combine GO enrichment in the context of gene pairs to compute functional similarities between two genes.

For a given pair of genes, term probabilities *p*(*t*) are evaluated using GOA data and *n*=2 and *k*= 1, or 2 are used for calculation of enriched term probabilities *p*\*(*t*).

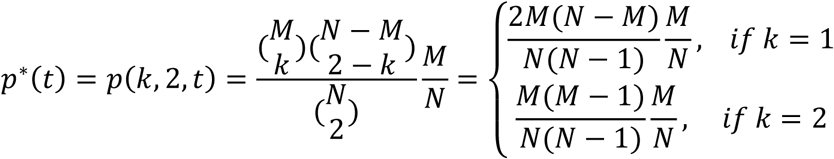

Enriched functional similarity *funsum*\* is obtained by *funsim* measures computed from enriched probability term *p*\*(*t*) instead of *p*(*t*) in computing information contents and semantic similarities.

## Acknowledgements

This work was partially supported by MOE2016-T2-1-029 AcRF Tier 2 grant by the Ministry of Education, Singapore.

